# *F_ST_* between Archaic and Present-Day Samples

**DOI:** 10.1101/362053

**Authors:** Diego Ortega-Del Vecchyo, Montgomery Slatkin

## Abstract

The increasing abundance of DNA sequences obtained from fossils calls for new population genetics theory that takes account of both the temporal and spatial separation of samples. Here we exploit the relationship between Wright’s *F_ST_* and average coalescence times to develop an analytic theory describing how *F_ST_* depends on both the distance and time separating pairs of sampled genomes. We apply this theory to several simple models of population history. If there is a time series of samples, partial population replacement creates a discontinuity in pairwise *F_ST_* values. The magnitude of the discontinuity depends on the extent of replacement. In stepping-stone models, pairwise *F_ST_* values between archaic and present-day samples reflect both the spatial and temporal separation. At long distances, an isolation by distance pattern dominates. At short distances, the time separation dominates. Analytic predictions fit patterns generated by simulations. We illustrate our results with applications to archaic samples from European human populations. We compare present-day samples with a pair of archaic samples taken before and after a replacement event.

---

Genomic sequences obtained from fossils provide new information about the history of present-day species. Already, thousands of partial or complete genomic sequences have been obtained from modern humans and their extinct relatives, and DNA sequences from fossils of numerous other species have been obtained as well (Reich 2018).

Population genetics theory of ancient DNA (aDNA) has focused primarily on the time dimension. Several methods have been developed to test for natural selection and estimate selection coefficients in a time series of samples (Bollback, et al. 2008; Malaspinas, et al. 2012; Terhorst, et al. 2015; Schraiber, et al. 2016). Much less effort has gone into incorporating the spatial dimension. The usual approach to analyzing spatially distributed aDNA is to use methods such as principal components analysis (PCA) and f-statistics that were developed for contemporaneous populations and ignore the ages of the fossils from which sequences are obtained. (Slatkin 2016)

There are three papers that have considered the spatial and temporal components of aDNA together. Skoglund et al. (2014) developed the coalescent theory of samples of different age and showed that PCA analysis can reveal the time separation of spatially distributed samples. Duforet-Frebourg and Slatkin (2016) extended the classic Kimura-Weiss (1964) analysis of isolation by distance in a stepping-stone model to predict the decrease in identity by descent with increasing spatial and temporal separation of samples. Silva et al. (2017) carried out an extensive simulation study that showed the importance of considering geographic structure when testing for population continuity. Although all these papers provide some insight into the effects of isolation by distance and time, they did not fully explore the effect on measures of population differentiation.

In this paper, we examine the effects of isolation by distance and time on pairwise *F_ST_* values. *F_ST_* and related statistics have been widely used to characterize isolation by distance. Using the principles introduced by Skoglund et al. (2014) and Duforet-Frebourg and Slatkin (2016), we will show how pairwise *F_ST_* between archaic and present-day samples reflects both the distance and time separating samples in equilibrium populations and in non-equilibrium populations after a partial population replacement.

## Pairwise *F_ST_*

*F_ST_* is useful for characterizing the extent of genetic difference between pairs of populations because it can be predicted analytically for a wide variety of models of population structure. If the per-locus mutation rate is small, *F_ST_* computed for pairs of populations is dependent on the average coalescence time of two copies of a gene, one drawn from each population (Slatkin 1991). We consider two populations *a* and *b*. We will use the Hudson et al (1992) estimator of *F_ST_*, which Bhatia et al. (2013) have shown has somewhat better properties than either the Weir and Cockerham (1984) or Nei (1986) estimators when applied to genomic data. Hudson et al. (1992) estimated *F_ST_* from the expression

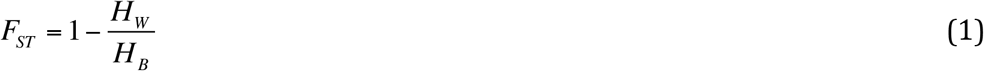

where *H_W_* is the average number of differences between chromosomes sampled from the same population and *H_B_* is the average number between different populations. That is, *H_W_* is the average of the expected per site heterozygosity within each population, which for two populations is 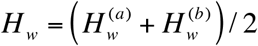 is the expected between-population heterozygosity.

Using the same method as in Slatkin (1991), we can find the expectations of *H_W_* and *H_B_* in terms of average branch lengths and the time between the samples when the per site mutation rate, *μ*, is small. For two lineages sampled at the same time, the average branch length is twice the average coalescence time. Therefore 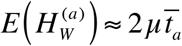 and 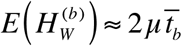 where *E* denotes the expectation and *t* ̅_*a*_ and *t*̅ _*b*_ are the average coalescence times of two copies of the locus sampled from populations *a* and *b*. Therefore 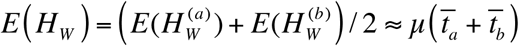.

If samples *a* and *b* are from different times, then no coalescence is possible until the lineage from the more recent sample reaches the time horizon of the older sample (Skoglund et al., 2014). Assume *a* and *b* were sampled *T_a_* and *T_b_* generations in the past, with *T_a_* < *T_b_*. Then *E* (*H_B_*) ≈ *μ*(*T_b_* – *T_a_* + 2*t*̅ *_ab_*), where *t*̅ _*ab*_ is the average coalescence time of the *a* and *b* lineages starting at *T_b_*. Therefore the expectation of the Hudson et al. estimator of *F_ST_* is approximately

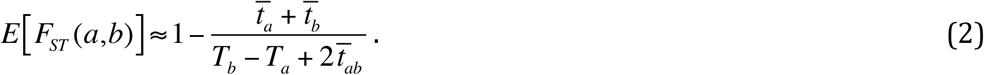

In many of the models, *t*̅_*a*_and *t*̅_*b*_ are the same for all populations while *t*̅_*ab*_ depends on the spatial separation of *a* and *b*.

It will be convenient to describe patterns of pairwise *F_ST_* in terms of the ratio

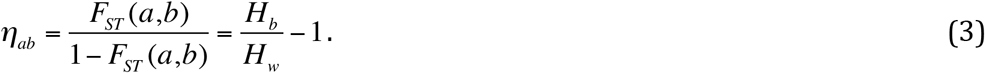

This ratio was introduced by Rousset (1997) and denoted by *β* / (1 – *β*). The Users Manual of Arlequin (Excoffier and Lischer 2010) called this ratio “linearized *F_ST_*”. From Eq. (2), it follows that

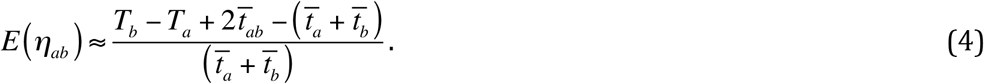

Thus, *η_ab_* is proportional to the additional average coalescence time between gene copies drawn from different populations attributable to their separation in space and time.

## Isolation by time

To illustrate our method, we consider first a single randomly mating diploid population. If the two samples come from the same population, the effect on *F_ST_* is easy to calculate. First, assume the population has constant effective size. Standard coalescent theory tells us *t* ̅_*a*_= *t*̅_*b*_= *t*̅ _*ab*_= 2*N*. Therefore 
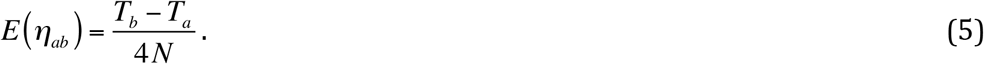

(Skoglund, et al. 2014). We compared the analytical estimates of *η_ab_* from Equation (5) with simulations in Supplementary Figure S2.

If the population size is a function of time, the result is not quite as simple. Both *t*̅_*a*_ and *t*̅ _*b*_ can be computed for an arbitrary demographic model from

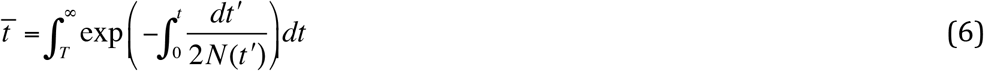

where *T*=*T_a_* or *T_b_*, and *t*̅_*ab*_= *t*̅ _*b*_. Therefore

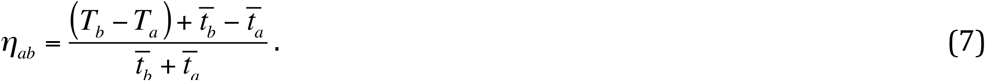

We can also obtain analytic results if samples are taken from sister populations. For simplicity, assume all populations are of effective size *N* and let the time of population divergence be *T_C_*. The two lineages cannot coalesce until they are in the ancestral population. Therefore, *t* ̅ *_a_* = *t*̅ _*b*_= 2*N* and *t*̅ *ab* = (*T_C_* – *Tb*) + 2*N* (Skoglund, et al. 2014) and

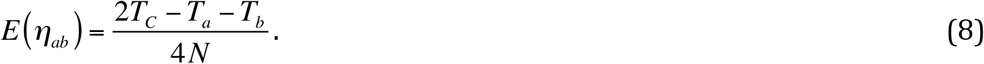

Thus, *E* (*η_ab_*) is proportional to the sum of the branch lengths in the population tree connecting the two samples. If population sizes depend on time in either or both branches, *η_ab_* would reflect the coalescence probabilities in the two branches. We compared the analytical estimates of *η_ab_* from Equation (8) with simulations in Supplementary Figures S1 and S3.

## Partial population mixture

We consider a generalization of a model analyzed by Skoglund et al. (2014) and illustrated by Figure 1. At time *t_C_* in the past the ancestral population splits into two descendent populations, *A* and *B*. The numbers in Figure 1 indicate the times of samples, with sample 1 being from the present day. At time *t_R_*, a fraction 1–*f* of population *A* is replaced by individuals from *B*. The resulting population continues to the present. How this model is described depends on the magnitude of *f*. If *f*=0, there was a complete population replacement. If *f* is small there was partial replacement. If *f* has an intermediate value, there was a population merger, and if *f* is nearly 1, there was admixture from *B* into *A*.

**Figure 1.**
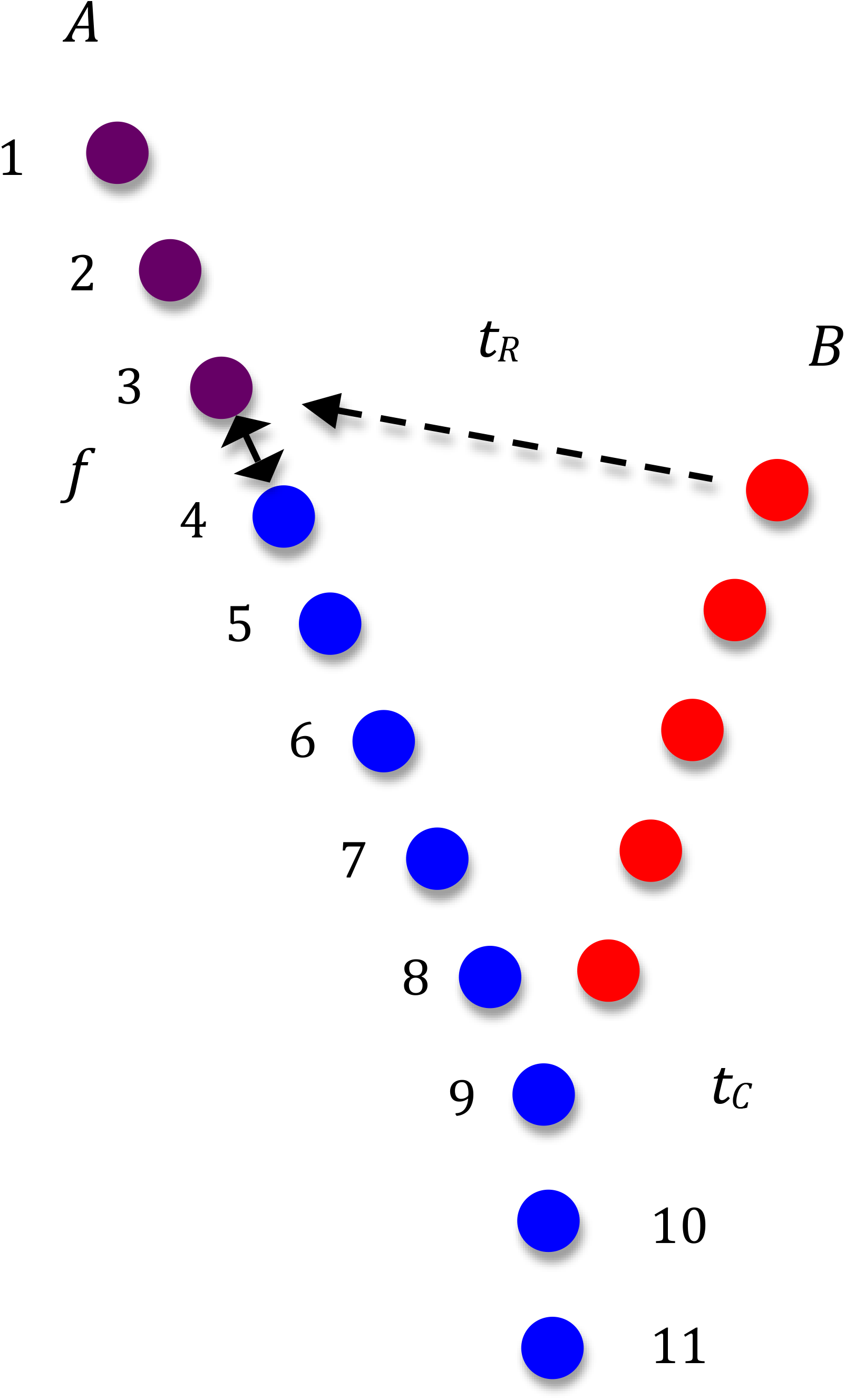
Illustration of the model of partial population replacement. This model is a generalization of one used by Skoglund et al. (2014). At time *t_C_* in the past, an ancestral population split into two descendent populations, distinguished as blue and red. Archaic samples are available from the blue population at different times in the past, indicated by the numbers. At time *t_R_* in the past, a fraction 1–*f* of the blue population is replaced by the red population. The resulting population survives to the present day.

To illustrate the main result, we assume all populations are of the same size, *N*. Variable population size can be accounted for in the same way as for a single population. We assume that the present-day population, sample 1, is compared to an archaic sample taken at time *τ* in the past. The average coalescence time *t*̅ of two lineages depends on whether the sample is taken before or after *t_R_*. If *τ* < *t_R_*, then

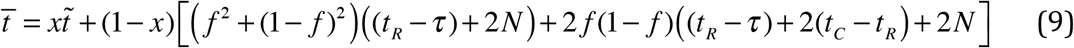

where 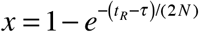 is the probability that the two lineages coalesce in the interval (*τ*, *t* _R_) and 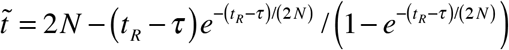 is the average time to coalescence given that they coalesce in that time interval. The logic is that if they coalesce before *t_R_*, the average coalescence time is *t* ̃. If they do not coalesce and both lineages go into the same ancestral population, the expected coalescence time is *t_R_* – *τ* + 2*N*. If they do not go into the same population, then they have to wait an additional *t_C_* – *t_R_* generations in each population before they can coalesce. If *τ* > *t_R_* then *t* ̅ = 2*N*.

We also need the between-sample coalescence time, *t*̅_*ab*_, If *τ* < *t_R_*, then

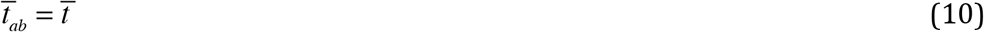

where *t*̅ is given by Equation (9). Once the present-day lineage reaches time *τ*, the average coalescence time is the same as if the two lineages were sampled at the same time. If *τ* > *t_R_*,

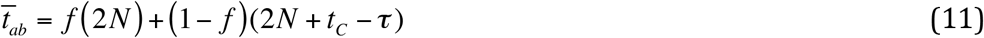
because with probability *f* the present day lineage remains in the same population and with probability 1–*f* it enters the other population. If *τ* > *t_C_*, *t*̅ _*ab*_= *τ / 2* + 2N.

Substituting these expressions into Equation (4), we can predict *η_ab_* as a function of *τ* and the other parameters. Some results are shown in Figure 2. The solid lines show the analytic predictions. The dots show simulation results obtained from using the program scrm (Staab, et al. 2015). In these and simulations described later in the paper, 100,000 replicates were run and results accumulated over all segregating sites. The mutation rate was chosen so that on average there were ten segregating sites per replicate. With this choice, there were no replicates with no segregating sites.

**Figure 2.**
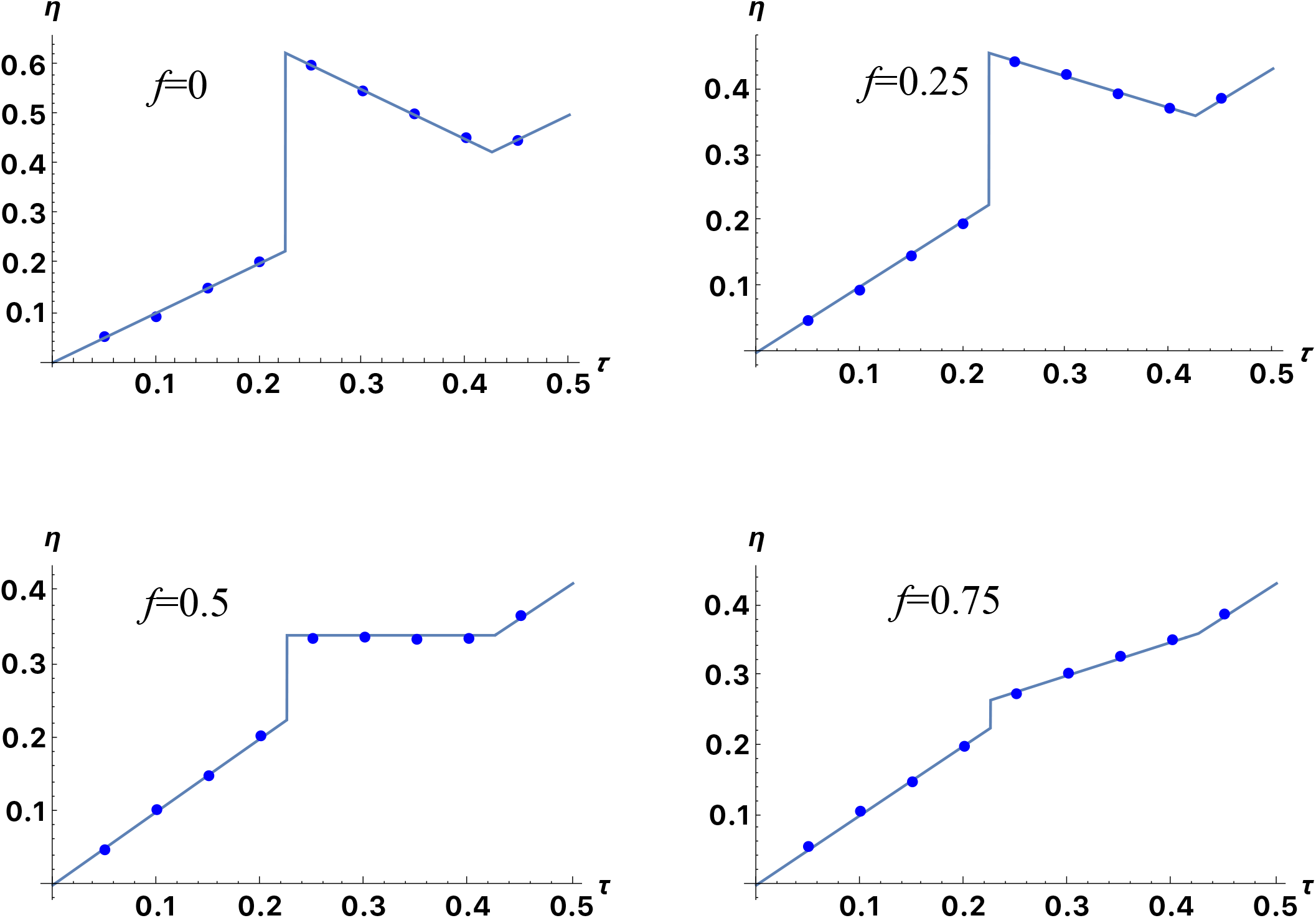
Comparison of analytic and simulation results quantifying the extent of differentiation (*η*) between a present-day population (1 in Fig. 1) and an archaic population (2 to 10 in Fig. 1) sampled before or after a partial population replacement. The analytic results indicated by the line were obtained from Equations (9)–(11) in the text. The simulation results indicated by the dots were obtained using scrm (Staab et al., 2015), where one chromosome was sampled from the present and the other sampled *τ* generations in the past, where *τ* is measured in units of 4*N* generations. The partial replacement occurred at 0.225(x4*N*) generations in the past.

## Isolation by distance and time

Duforet-Frebourg and Slatkin (2016) showed that the combined effects of isolation by distance and time in a stepping-stone model can be understood by considering the movement of lineages ancestral to the more recent sample during the time interval between the two samples. That movement is governed by dispersal patterns during the interval. Coalescence cannot occur until the time of the older sample. For simple models, analytic results can be obtained.

To illustrate, consider a one-dimensional stepping stone model with *d* demes arranged in a circle, and assume a migration rate *m* between adjacent demes. The average coalescence time of two genes sampled from *i* steps apart is 
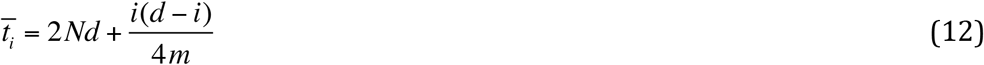
where here *i* is counted in a clockwise direction (0≤*i*≤*d*–1) (Slatkin 1991). To see the effect of the difference in sampling time, assume one sample is from the present, (*T_a_* = 0) and the other from *T* generations in the past (*T_b_* = *T*). Between 0 and *T*, the present-day lineage undergoes a random walk on the circle. The probability that the lineage will be in deme *j*, given that it was initially in deme *i* is *p_ij_*, the jth element of the vector, 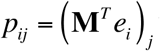 where *e_i_* is a unit vector with 1 in position *i* and 0 otherwise and **M** is a circular matrix which has non-zero elements *M_ii_* = 1 – 2*m* and *M_i,j+1_* = *M_i,j–1_* = *M _1,d_*= *M_d,1_* = *m*. **M**^*T*^denotes the Tth power of **M.**

Therefore *t*̅_*a*_= *t*̅_*b*_= 2*Nd* and

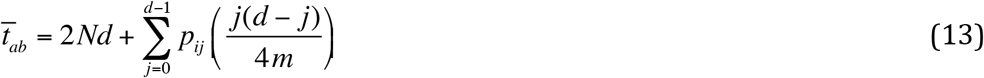

from which we can compute *η_i_* to be

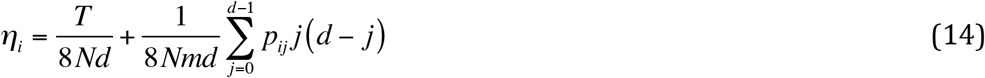

Figure 3 presents some typical results for the case with archaic samples drawn from deme 0 at various times. Shown for comparison is the equilibrium IBD pattern for contemporaneous samples (*T*=0). As the age of the archaic sample increases, *ηi* increases in the neighborhood of the sampled deme. There are two components to this increase. One is the time separation of the samples, represented by the first term in Equation (14). The other is the averaging of the equilibrium pattern because of the dispersal of the present-day lineage between 0 and *T*, which is represented by the second term in Equation (14). Because *ηi* = 0 for two samples from the same population at the same time, the averaging is over populations for which *ηi* is positive. That results in a positive contribution. Both terms contribute and their relative magnitudes depend on the parameter values. Note that in this model, as in many models of a subdivided population, the average within-deme coalescence time is twice the total number of individuals in the population, independently of the migration pattern (Strobeck 1987).

**Figure 3.**
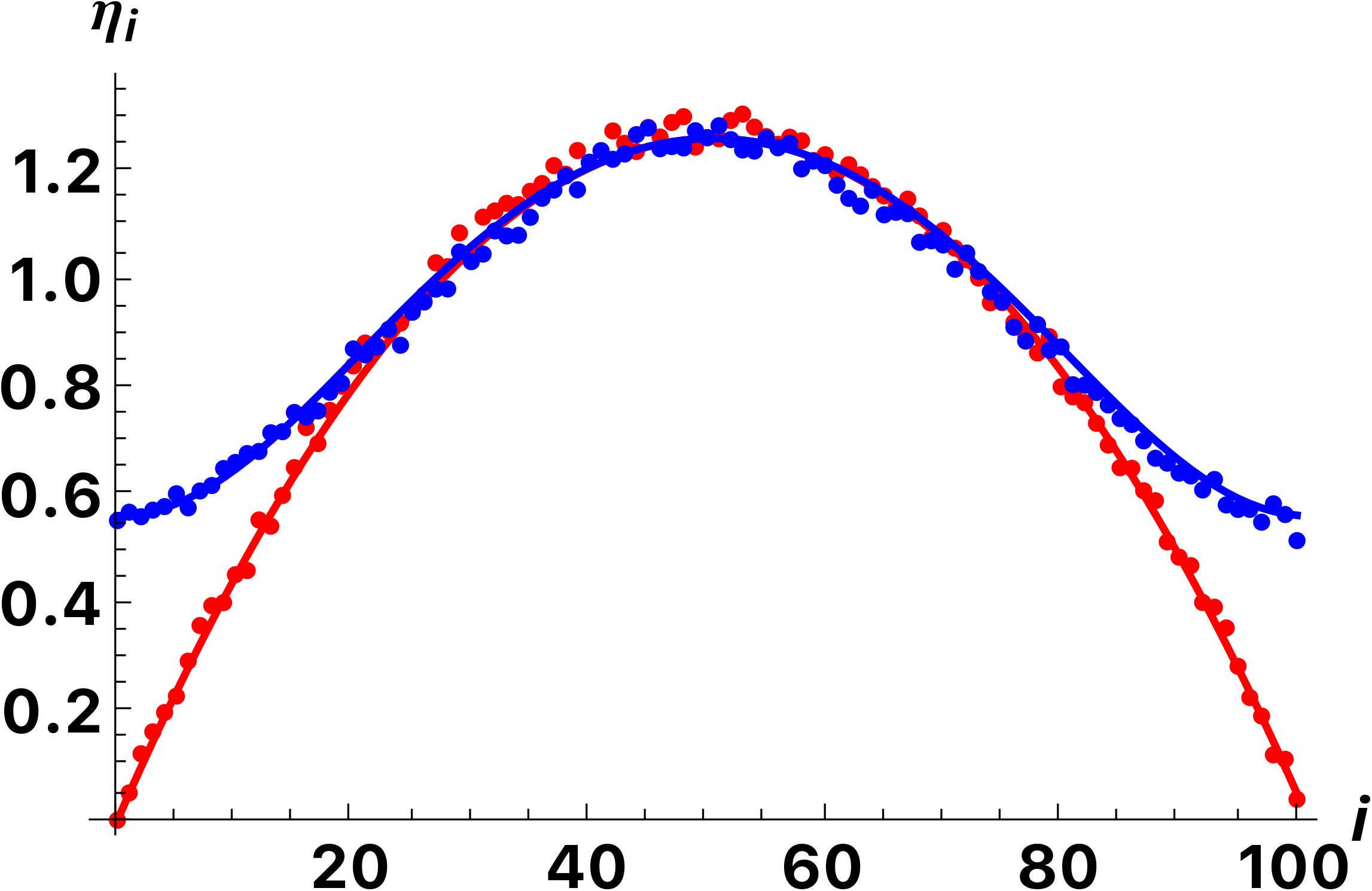
Comparison of analytic and simulation results quantifying the extent of differentiation (*ηi*) between populations *i* steps apart in a circular stepping-stone population sampled at the same time (red) or with a time-separation of 40*N* generations (blue). Comparison of analytic and simulation results for a circular stepping stone model. The analytic results, shown by the solid lines, were obtained using Equation (14) in the text. The simulation results, shown by the dots, were obtained using scrm (Staab et al., 2015). The model was of a circle of 101 demes with migration rate 4*Nm*=10 between adjacent demes. The red dots and line show the equilibrium pattern of isolation by distance between contemporaneous populations. The blue dots and line show the pattern for a sample taken 40*N* generations in the past. Simulation results are averages of 100,000 replicates.

Similar results are obtained for one and two dimensional symmetric stepping stone models. Figure 4 shows typical examples.

**Figure 4.**
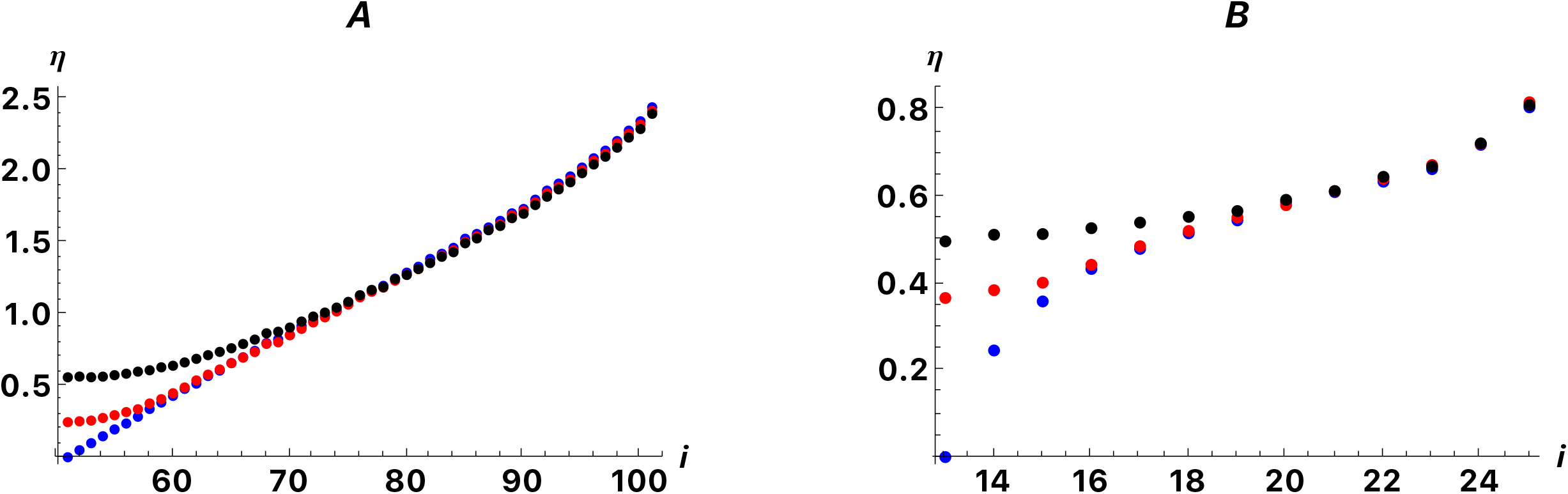
Isolation by distance patterns in one and two dimensional stepping stone models with symmetric migration. Pairwise values of *η* were estimated from simulation results obtained with scrm (Staab et al. 2015). Each point is based on 100,000 replicates. Part A. 101 × 1 stepping stone model with 4*Nm*=1 between adjacent demes. The middle population (population 51) was sampled at the present (*t*=0, Blue dots), and two times in the past, *t*=8*N* generations (Red dots) and *t*=40*N* generations in the past (Black dots) and compared to each of the present-day populations. The results are symmetric around the middle population. Part B. 25 × 25 stepping stone model with 4*Nm*=1 between adjacent demes. The middle population in the middle row (population 13,13) was sampled at the present (*t*=0, Blue dots), and two times in the past, *t*=8*N* generations (Red dots) and *t*=40*N* generations in the past (Black dots) and compared to each of the present-day populations in the middle row (population 1,13 to population 13,13). The results are symmetric around the middle population. Note the difference in scale of the vertical axes in Parts A and B.

Models of symmetric dispersal are a staple of population genetics theory because of their mathematical simplicity. There is no reason to suppose that dispersal in natural populations is actually symmetric either in each generation or when averaged over many generations. Comparison with archaic samples can reveal slight asymmetry in dispersal that may not be apparent when comparing only present-day samples. Figure 5 provides one example. The population is in a 101×1 linear stepping stone model, as in Figure 4A. The only difference is that 4*Nm* to the right and left are 11 and 9 respectively. As shown in Fig. 5, this difference is not obvious in the isolation by distance pattern of present day populations, but is when a few archaic samples are included.

**Figure 5.**
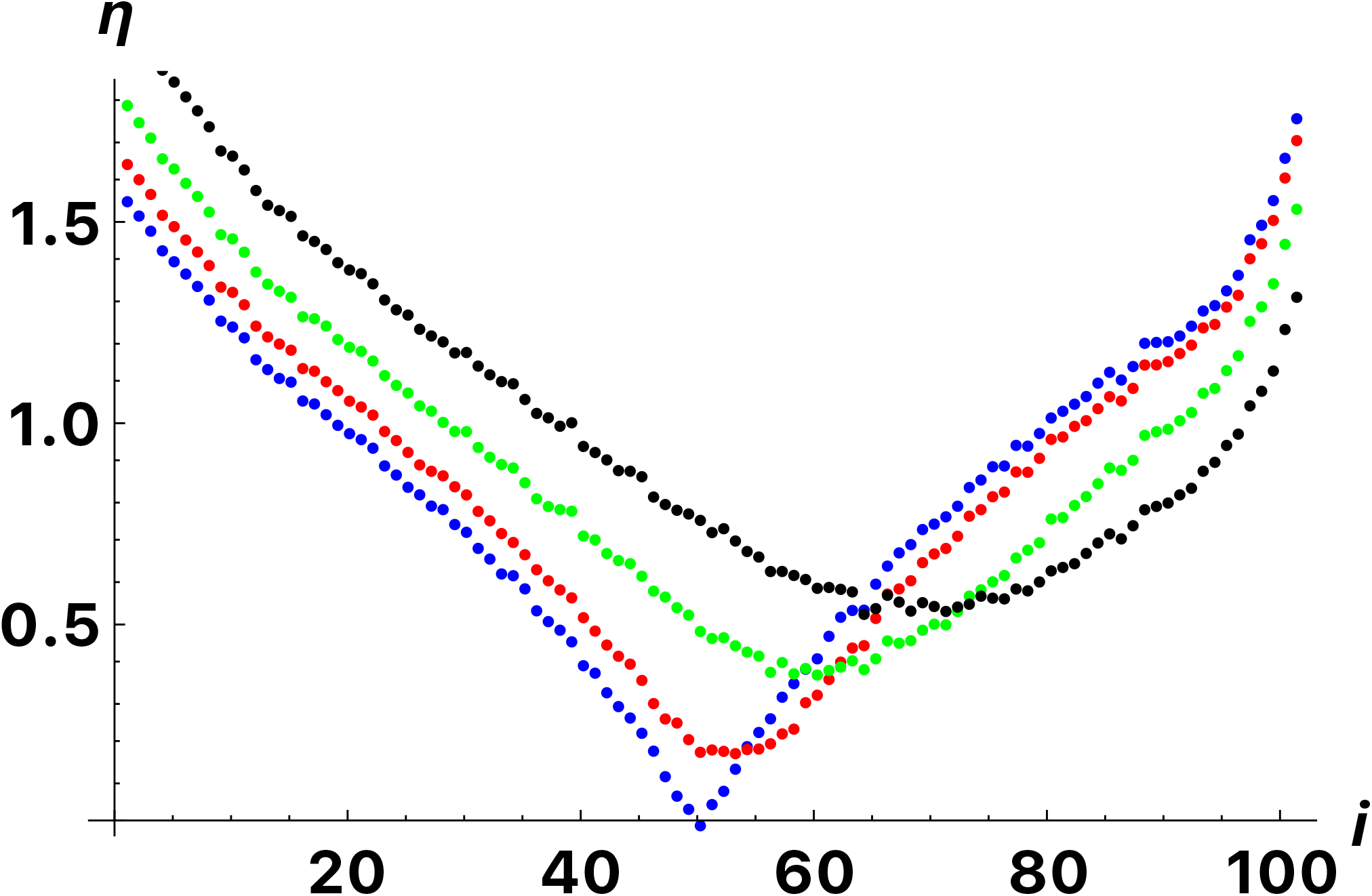
Simulation results for a 101×1 stepping stone model with asymmetric migration. The model is the same as in part A of Figure 4 but with migration rate to the right of 4*Nm*=11 and to the left of 4*Nm*=9. We show values of *η* for archaic samples taken at 4*N* (red), 8*N* (green) and 12*N* (black) generations in the past compared to each present-day population. The blue dots are for a sample taken at *t*=0.

## Range expansion with partial replacement

Range expansions have happened many times in the history of humans and other species. Range expansions create unusual patterns in allele frequencies because of continued founder effects, a phenomenon called “gene surfing”. (Excoffier and Ray 2008) Several ways have been proposed for detecting the genetic signatures of range expansions including testing for clines in heterozygosity (Ramachandran, et al. 2005) and computing a directionality index (Peter and Slatkin 2013). Range expansion may occur into an area previous unoccupied or an area occupied by another population. In the latter case, admixture between the invading and resident populations will take place. (Currat, et al. 2008).

To determine the effects of range expansion on *F_ST_* taken from archaic samples, we simulated a model in which there is a partial replacement of a resident population by an expanding population. Both before and after the range expansion, there is a stepping stone population structure. The model is illustrated in Figure 6. As in Figure 1, *f* is the fraction of each population that is descended from the resident population and 1–*f* is the fraction that is descended from the invading population in each location. Some simulation results are shown in Figures 7 and 8. The patterns in pairwise *η* values are a combination of those seen with partial population replacement and isolation by distance in an equilibrium population. The pattern of isolation by distance and the relationship between archaic and present-day samples are preserved if there is partial replacement (Figure 7). And the abrupt change created by a partial replacement is evident when comparing archaic samples before and after the replacement event (Figure 8).

**Figure 6.**
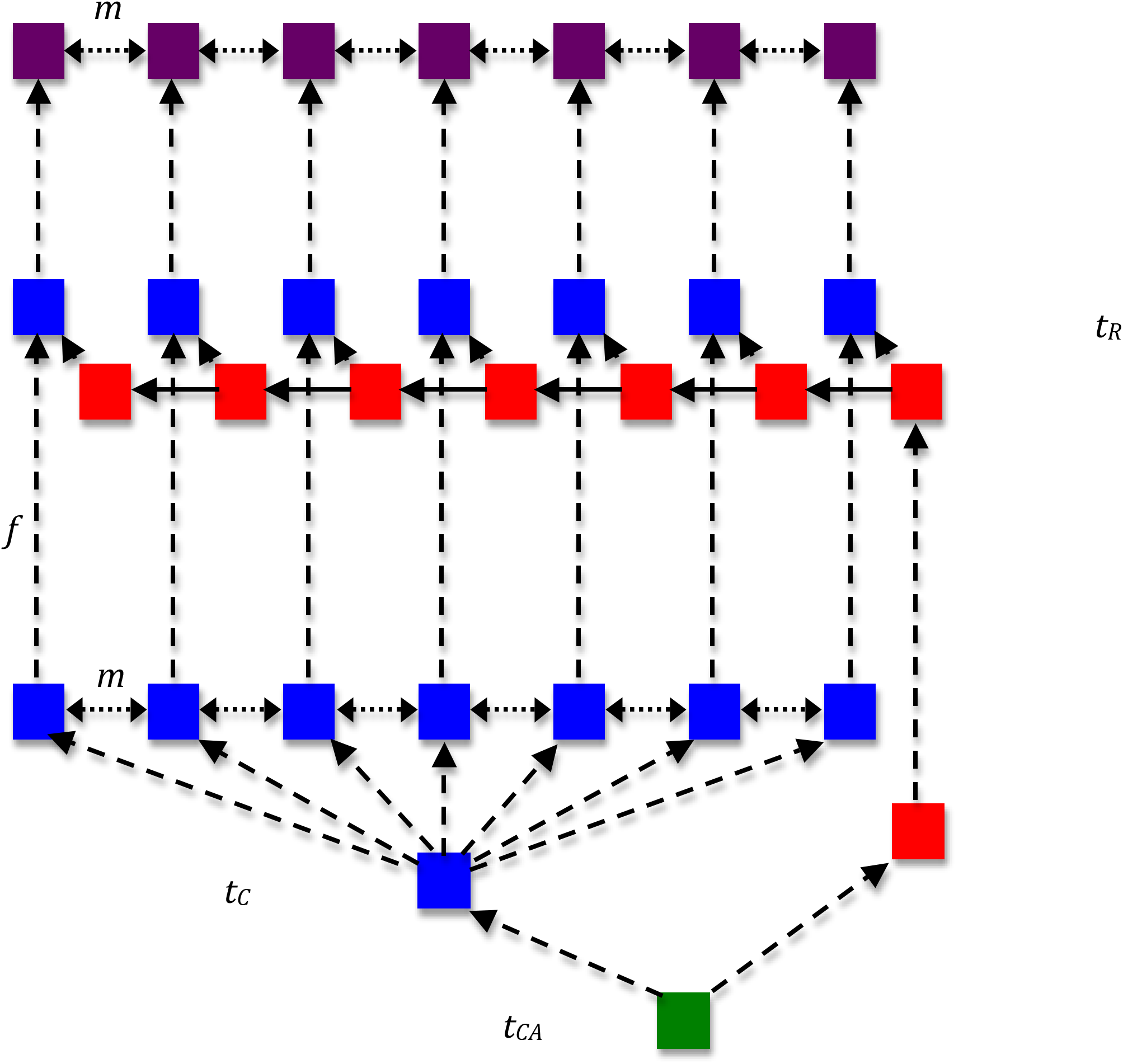
Illustration of the model of partial population replacement in a stepping stone framework. At time *t_CA_* in the past, the ancestral (green) population split into two descendent populations, blue and red. At time *t_C_*, the blue population gave rise to several populations which then exchange migrants at rate *m* in a stepping stone model. At time *t_R_*, the descendant of the red population becomes the source for a range expansion. After each colonization event, the red population mixes with a blue population to produce the purple descendent population. Each descendent population is made up of a fraction *f* of the resident (blue) population and a fraction 1–*f* of the expanding (red) population. The purple populations exchange migrants symmetrically with each neighboring population at rate *m* until the present day.

**Figure 7.**
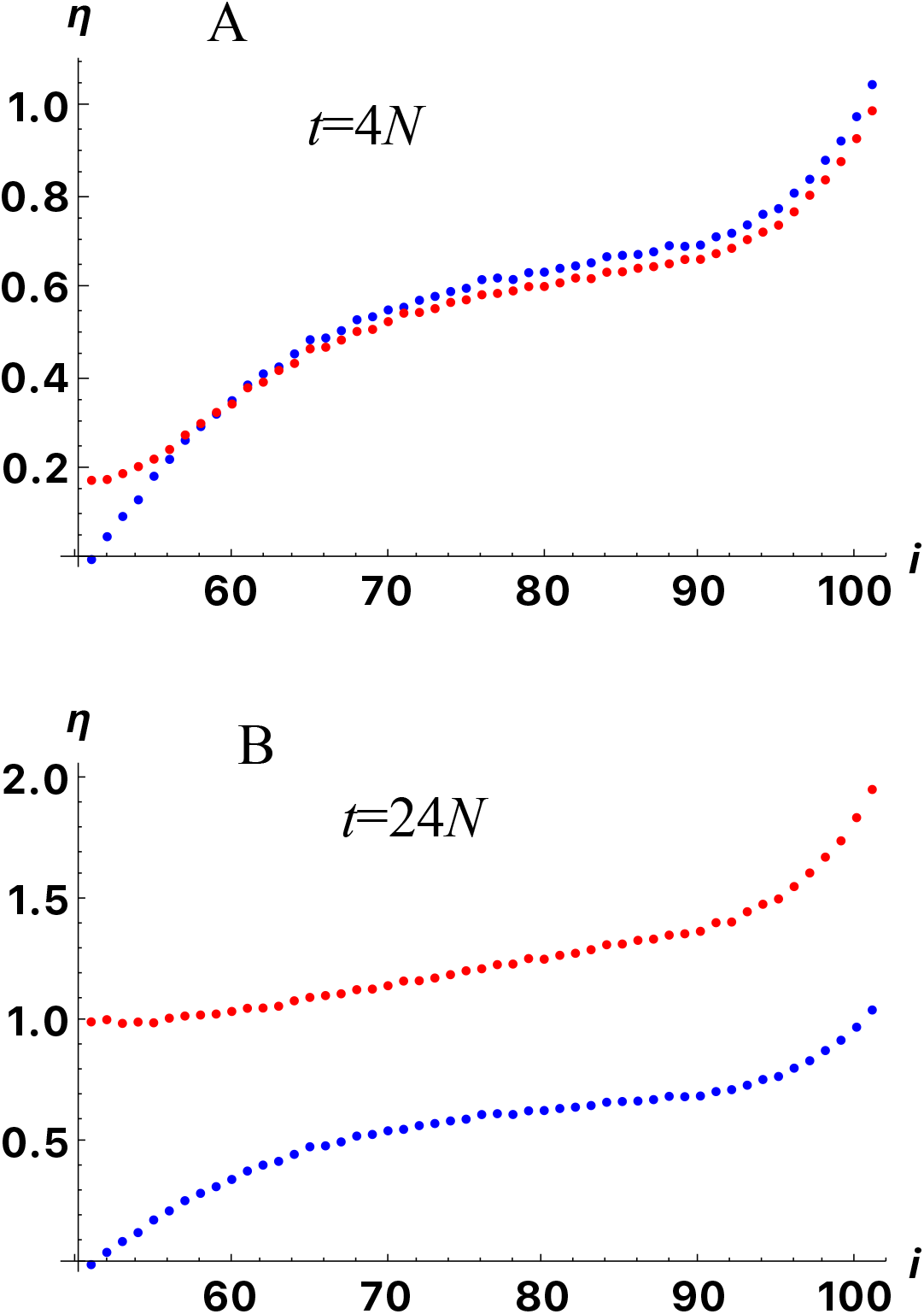
Representative patterns of isolation by distance seen when archaic samples are taken before and after a partial replacement. The model is as illustrated in Figure 6. There were 101 populations in a stepping stone configuration with migration at rate 4*Nm*=10 between adjacent demes. At time *t*=16*N* in the past, there was a range expansion beginning with population 1. During each colonization event, the population size was reduced by a factor of 0.01 for 0.002N generations. As each colonizing population came into contact with a resident population, the two populations contributed equally to the descendent population (*f*=0.5). The blue dots indicate *η_ab_* for the middle present-day population compared to each of the other present-day populations. The red dots indicate the values for the middle archaic population and each of the present-day populations. The two archaic samples were taken *t*=4*N* and *t*=24*N* generations before the present. The graphs were obtained using scrm. Each point is the average of 100,000 replicate simulations.

**Figure 8.**
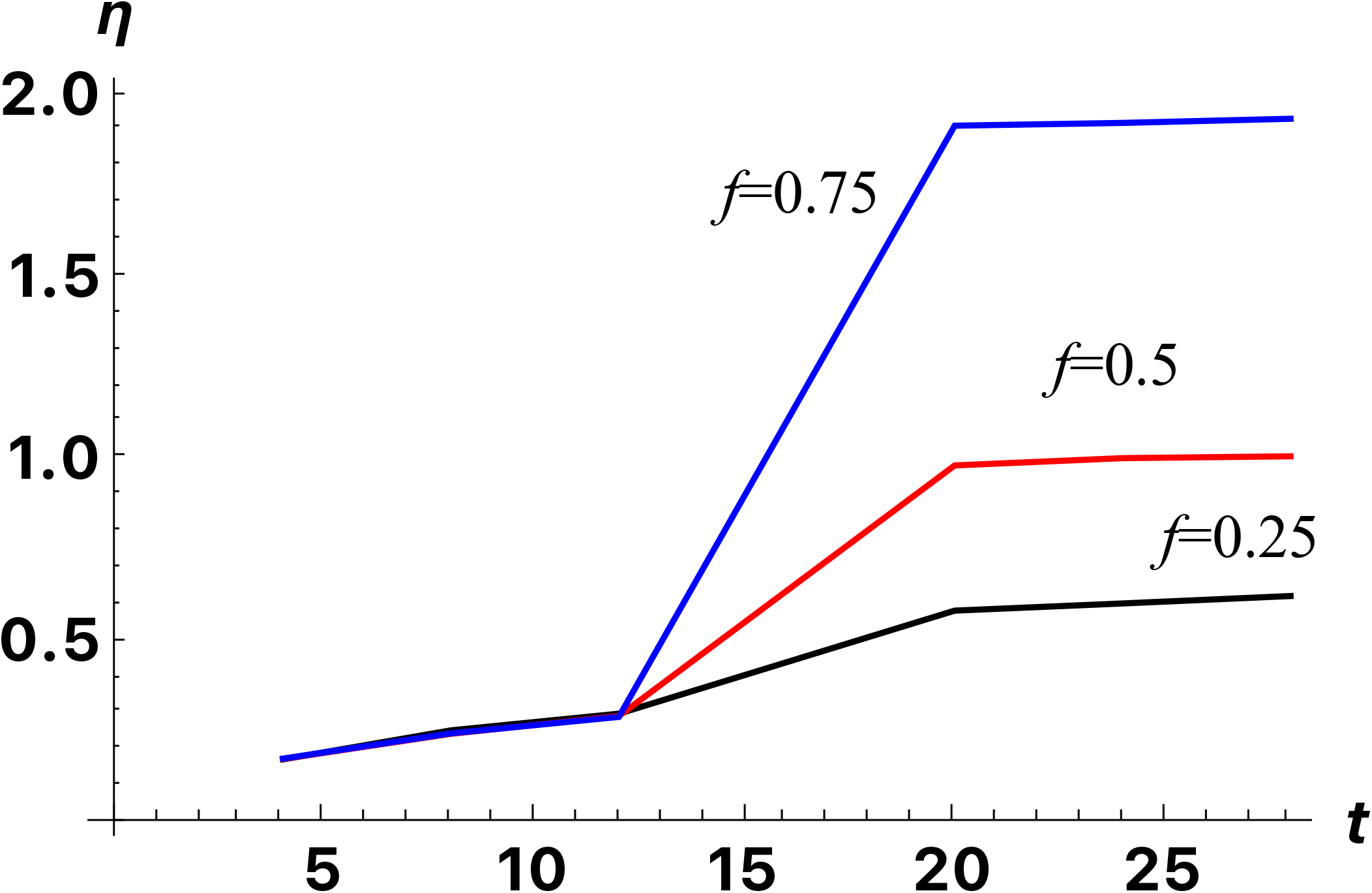
Comparison of increases in *η* for different extents of admixture with resident populations. The parameters were the same as in Figure 7 except that the results for *f*=0.25, 0.5 and 0.75 are shown. The archaic samples were taken 4*N*, 8*N*, 12*N*, 20*N*, 24*N* and 28*N* generations in the past.

## Examples

To illustrate patterns seen in data from human populations, we reanalyzed the data of the Simons Genome Diversity Project (Mallick, et al. 2016) and two ancient human genomes (Lazaridis, et al. 2014). The ancient genomes come from a Neolithic farmer (the Stuttgart sample, ~7000 years before present) and a Neolithic hunter-gatherer (the Loschbour sample, ~8000 ybp). Figure 9 shows the results. The red histograms in Fig. 9 show pairwise values of *η_ab_* computed for Stuttgart and several present-day European samples. The orange histograms show *η_ab_* computed for Loschbour and the same present-day samples. The results are consistent with two theoretical expectations: The older sample, Loschbour, has larger *η_ab_* values. Additionally, the results are consistent with a smaller average ancestry in present-day Europeans coming from hunter-gatherers (Haak, et al. 2015). This is in agreement with our partial population replacement model, where comparisons of present-day individuals with ancient samples coming from a population that has been mostly replaced (*f* close to 0) tend to have larger *η_ab_* when the ancient sample was sampled from before the time of replacement, *t_R_*, and after the present-day and ancient populations coalesce to an ancestral population, *t_c_*, (see Figure 2). The ancient samples we used, Loschbour and Stuttgart, are samples taken from near the time of replacement.

**Figure 9.**
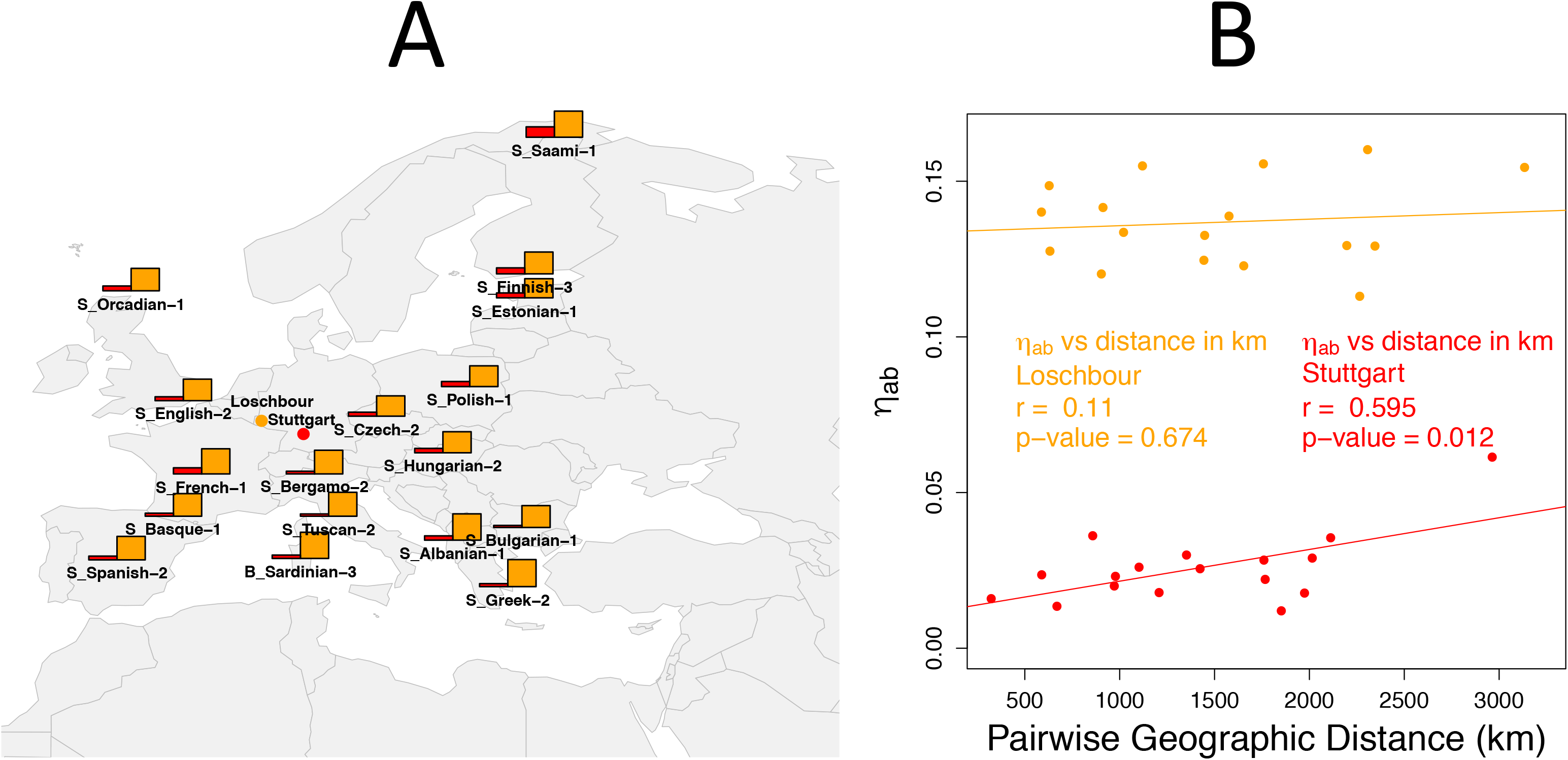
A. Comparison of pairwise *η_ab_* values computed for two focal samples. All data were taken from the Simons Genome Diversity Project dataset (Mallick et al., 2016), which also contains two ancient human genomes, Stuttgart and Loeschbour (Lazaridis et al., 2014). We compare the results for the two focal samples. The red dot indicates the location of the Stuttgart Neolithic farmer skeleton (~7,000 years old) and the orange dot points the location of the Loschbour Neolithic hunter-gatherer skeleton (~ 8,000 years old). The histogram bars indicate the value of *η_ab_* computed between the focal sample of the same color and a present-day sample at each location. B. Pairwise *η_ab_* values of present-day samples and the two focal ancient samples vs. the pairwise geographical distance between the sampling location of the present-day and ancient samples. The correlation coefficient *r* and the p-value of the null hypothesis that the slope obtained from the linear regression line has a value different from zero. The p-values were obtained using an F-test comparing the linear model with a non-zero slope to a model with a zero slope.

We found a significant positive correlation between the pairwise geographical distance and the pairwise *η_ab_* values of present-day samples and the Stuttgart sample (Figure 9B). This observation is consistent with a pattern of isolation by distance in Neolithic farmers that is retained in present-day populations. In contrast, there is no significant correlation when we do the same analysis with the Loschbour sample. This observation suggests that the replacement of hunter-gatherer populations by early farmers erased any signal of isolation-by-distance in the hunter-gatherer populations, if one was present.

## Discussion and Conclusion

We present the basic theory of Wright’s *F_ST_* between samples taken at different times and places. As Skoglund et al. (2014) note, there is an important difference between pairwise *F_ST_* and the principal components analysis (PCA). Pairwise *F_ST_* values do not depend on what other samples are included in the analysis while principal components do. Although both representations of data reflect pairwise coalescence times (Slatkin 1991; McVean 2009), principal components depend on pairwise coalescence times for a particular pair of samples relative to other pairs of samples. The two ways of looking at data are both useful. Using pairwise *F_ST_* values allows a more direct tie to the underlying coalescent process and allows comparison with analytic theory.

The theory we have developed shows that *F_ST_* values for pairs of samples of different age depend on numerous parameters in addition to the time separation of the samples. For that reason, pairwise *F_ST_* values alone are not suitable for inferring demographic parameters. Both the results presented here and the simulation study of Silva et al. (2017) show that patterns of population differentiation depend in a complex way on the time separation of samples, patterns of dispersal and the extent of population replacement. However, pairwise *F_ST_* values could serve a key statistics in an approximate Bayesian computation analysis (Bertorelle, et al. 2010) because they directly reflect pairwise coalescence times.

## Acknowledgements

This research was supported in part by a US NIH grant, R01-GM40282 to M. S. We thank L. Excoffier for helpful comments on an earlier version of this paper.

---

**Supplementary Figure S1.**
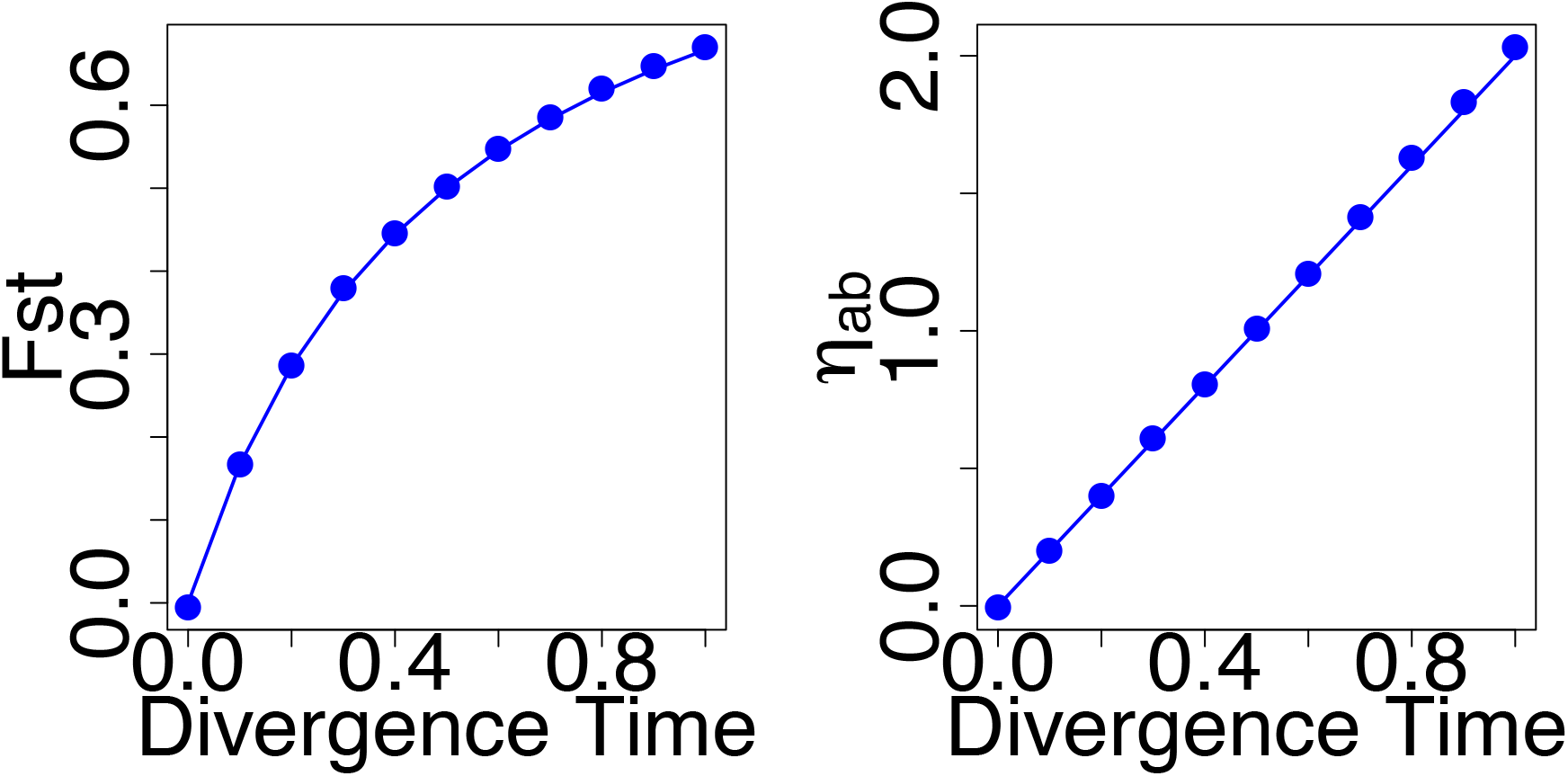
Estimates of *Fst* and *η_ab_* in a demographic model where two demes diverged from an ancestral deme without subsequent migration. The ancestral deme and the two present day demes have a population size N= 10,000. The divergence time, shown in the x-axis in both plots, is measured in units of *4N* generations. The dots in the plots represent the average values of *F_st_* and *η_ab_* across 100,000 independent sites from simulations done using *ms* (Hudson (2002), Bioinformatics), where two chromosomes were sampled from each of the two populations. The lines show the analytical results, using *t_a_* = *t_b_* = 20000 and *t_ab_* = 20000 + *D* * 40000, where *D* is equal to the divergence time measured in units of *4N* generations. We used Equations (2) and (8) for the analytical results of the left and right plot, respectively.

**Supplementary Figure S2.**
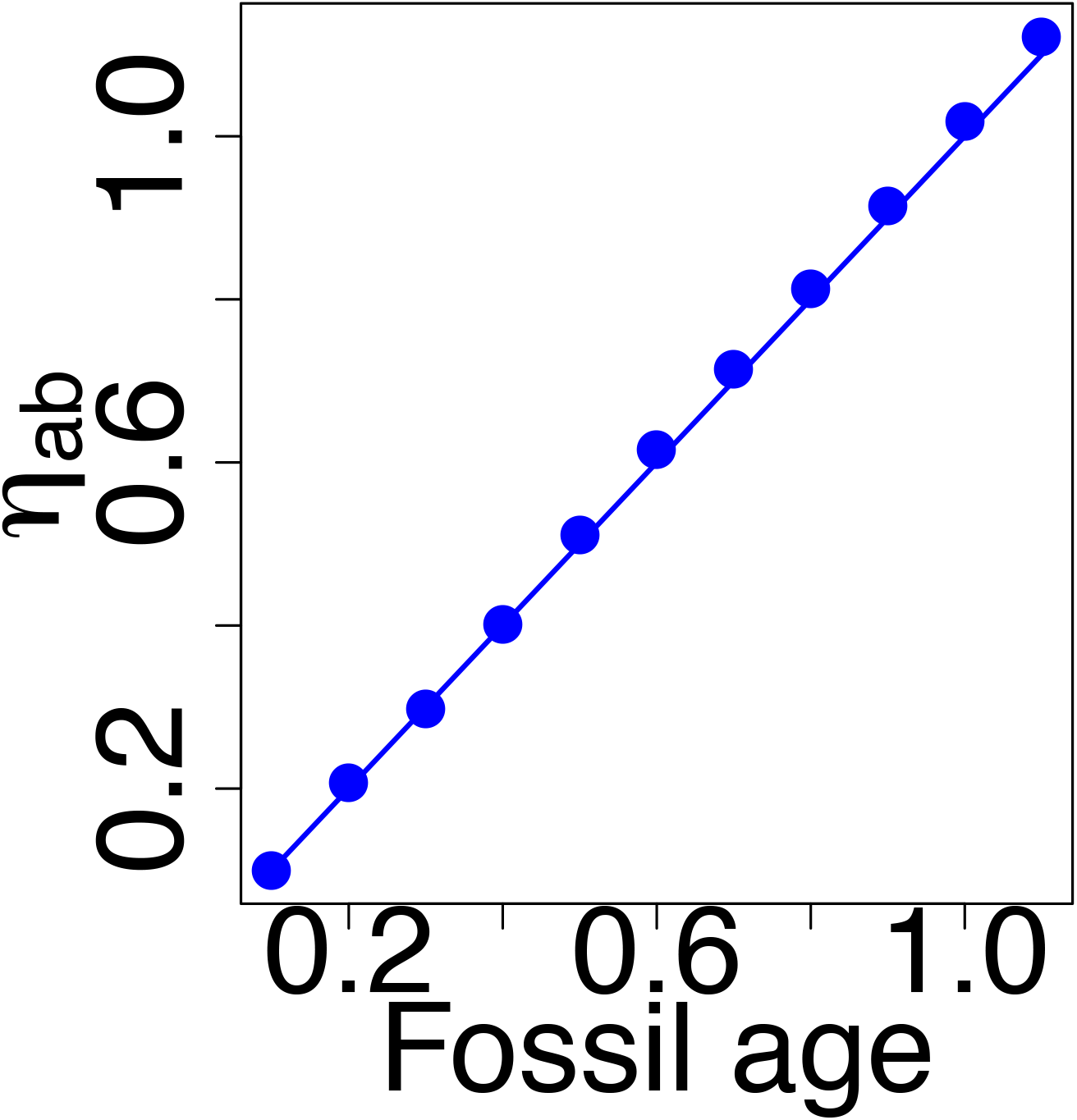
Estimates of *η*_ab_ in a demographic model with one population. The calculations of *η*_ab_ performed here were done sampling two chromosomes from the present (*T_a_* = 0) and the other two chromosomes were taken from a fossil. The fossil age is measured in units of 4*N* generations and is shown in the x-axis. The deme has a population size N = 10,000. The dots in the plots represent the average value of η_ab_ across 100,000 independent sites from simulations done using *ms* (Hudson (2002), Bioinformatics). The lines show the analytical results, using *N* = 10000, *T_a_* = 0.0 and *T_b_* = *F* * 40000, where *F* is the fossil age measured in units of *4N* generations. We used Equation (5) to obtain the analytical results.

**Supplementary Figure S3.**
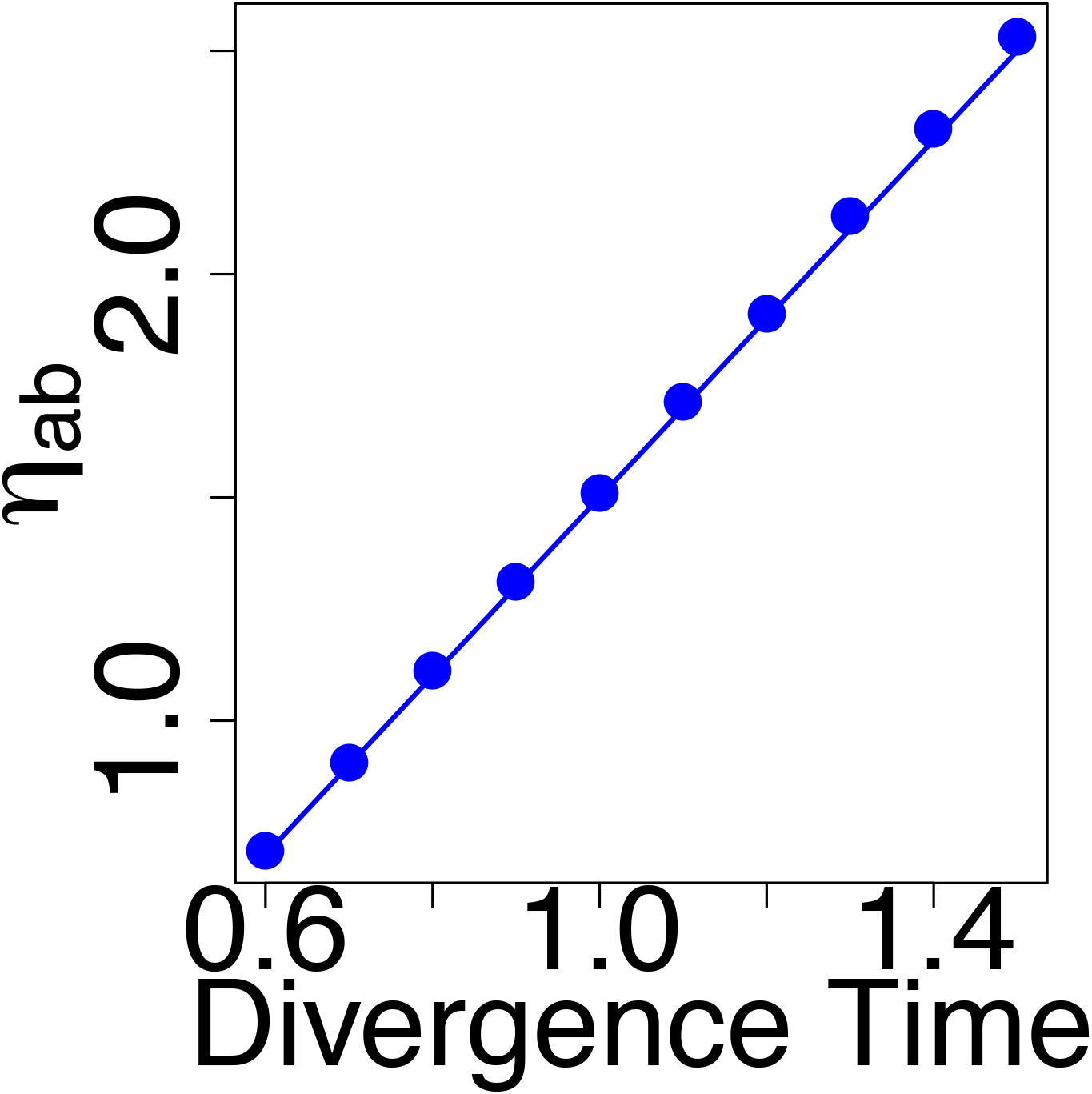
Estimates of η_ab_ in a demographic model where two demes diverged from an ancestral deme without subsequent migration. Two chromosomes were sampled from the first deme at time *T_a_* = 0. Another two chromosomes were sampled from a fossil at time *T_b_* = 20,000. The ancestral deme and the two present day demes have a population size N= 10,000. The divergence time, shown in the x-axis, is measured in units of *4N* generations. The dots in the plots represent the average value of *η_ab_* across 100,000 independent sites from simulations done using *ms* (Hudson (2002), Bioinformatics). The lines show the analytical results, using *T_c_* = *D* * 40000, where *D* is equal to the divergence time measured in units of *4N* generations. We used Equation (8) to get the analytical results.

